# Structured clustering of the glycosphingolipid GM1 is required for membrane curvature induced by cholera toxin

**DOI:** 10.1101/2020.01.22.915249

**Authors:** Abir Maarouf Kabbani, Krishnan Raghunathan, Wayne I. Lencer, Anne K. Kenworthy, Christopher V. Kelly

## Abstract

AB_5_ bacterial toxins and polyomaviruses induce membrane curvature as a mechanism to facilitate their entry into host cells. How membrane bending is accomplished is not yet fully understood but has been linked to the simultaneous binding of the pentameric B-subunit to multiple copies of their glycosphingolipid receptors. Here, we probe the toxin membrane binding and internalization mechanisms by using a combination of super-resolution and polarized localization microscopy. We show that cholera toxin subunit B (CTxB) can induce membrane curvature only when bound to multiple copies of its glycosphingolipid receptor, GM1, and the ceramide structure of GM1 is likely not a determinant of this activity as assessed in model membranes. A mutant CTxB capable of binding only a single GM1 fails to generate curvature either in model membranes or in cells and clustering the mutant CTxB-single-GM1 complexes by antibody cross-linking does not rescue the membrane curvature phenotype. We conclude that both the multiplicity and specific geometry of GM1 binding sites are necessary for the induction of membrane curvature. We expect this to be a general rule of membrane behavior for all AB_5_ toxins and polyomaviruses that bind glycosphingolipids to invade host cells.

**SIGNIFICANCE STATEMENT:** Membrane binding toxins demonstrate both a public health challenge and a bioengineering opportunity due to their efficient internalization into cells. These toxins multivalently bind to naturally occurring lipid receptors at the plasma membrane and initiate endocytosis. This manuscript reports the importance of structured lipid-receptor clustering for the induction of membrane bending. We also observed that the magnitude of membrane curvature was correlated to the stoichiometry of toxin-bound receptors. By identifying how these bacterial proteins initiate membrane curvature, these findings provide mechanistic insights into the early steps of pathogenic endocytosis.

## INTRODUCTION

Cholera toxin (CTx) causes the massive and often deadly secretory diarrhea following enteric infection with the bacteria *Vibrio cholerae.* 2.9 million infections and 95,000 deaths are attributed to cholera annually (1). CTx typifies the AB_5_ family of bacterial toxins that includes the heat-labile *E. coli,* Shiga toxin (STx), tetanus, and pertussis toxins. The B-subunits of these toxins multivalently bind and cross-link naturally occurring plasma membrane glycosphingolipids to induce cellular internalization and guide intracellular trafficking (2). CTx, STx, and simian virus 40 (SV40) are known to induce membrane bending, likely through binding to multiple copies of their glycolipid receptors with acyl tail dependencies (3–9). In the case of CTx, the B-subunit (CTxB) binds the glycosphingolipid GM1 to engage endogenous mechanisms of endocytic uptake and retrograde trafficking from the plasma membrane into the endoplasmic reticulum (ER). Once in the ER, the enzymatically active A-subunit co-opts the machinery for ER-associated degradation to retro-translocate into the cytosol and induce disease (10). Except for pertussis toxin, the B-subunits of these toxins assemble into stable and perfectly symmetrical homopentamers that can bind five or more glycosphingolipids at once and localize the toxin within plasma membrane nanodomains enriched in cholesterol (11, 12). But, how these evolutionarily conserved and highly-potent B-subunits work to enable invasion of the host cell remains incompletely understood.

Here, we test the idea that CTxB requires defined stoichiometry of binding to induce *de novo* curvature in membranes (13–17) and to sort the toxin into such highly-curved membrane structures (17–19). Because CTxB internalization *in vivo* does not exclusively rely on either clathrin-or caveolin-dependent processes (20–27), it is feasible that the inherent membrane bending capability of CTxB may drive cellular uptake (3, 4). We examine the mechanisms by which CTxB induces membrane curvature using polarized localization microscopy (PLM). PLM provides super-resolution information on membrane orientation and detects membrane orientation with order-of-magnitude higher sensitivity than comparable optical methods (28). Curvature induction was measured in model membranes to test the influence of the multivalent binding of CTxB and the molecular shape and lipid phase preference of GM1. We found that membrane bending in model membranes caused by CTxB can be explained by the stoichiometry of binding at least two GM1 molecules per CTxB, and no contribution was observed from the ceramide tail structure of the GM1 or membrane cholesterol content. Finally, the need for multivalent binding of CTxB to GM1 to induce membrane curvature in the plasma membrane of live cells was confirmed by super-resolution imaging of cells exposed to pentavalent wild type (wt) or a monovalent mutant CTxB (mCTxB) capable of binding only a single GM1.

## RESULTS

### Stoichiometry of GM1 binding to CTxB dictates membrane curvature induction

We tested the hypothesis that AB_5_ toxins employ a structured cross-linking of their ganglioside receptor to induce membrane curvature. We tested this with CTxB that contains 5 potential binding sites for its receptor GM1 as a model system. To detect membrane curvature, we used a nanoscale membrane budding assay based on PLM (28). Using this assay, we previously showed the addition of fluorescently labeled CTxB to supported bilayers composed of POPC, DiI, and GM1 induced the growth of membrane buds with a radius of < 50 nm that subsequently grow to form tubules with a radius of > 200 nm. These membrane bending events were driven by the accumulation of CTxB at the base of the tubule, reflecting the preference of the toxin for negative Gaussian membrane curvature vs. planar or positive-curvature regions.

We first tested the effect of receptor clustering by altering the average number of GM1s bound per toxin. This was accomplished by either varying the concentration of CTxB while holding the levels of GM1 in the membrane constant at 0.3 mol% or varying the concentration of GM1 while holding the CTxB concentration constant at 4.3 nM (Fig. 1). Membrane curvature and CTxB localization were then determined independently via p-polarized localization microscopy (pPLM) and direct stochastic optical reconstruction microscopy (dSTORM), respectively (28, 29).

**Figure 1:**
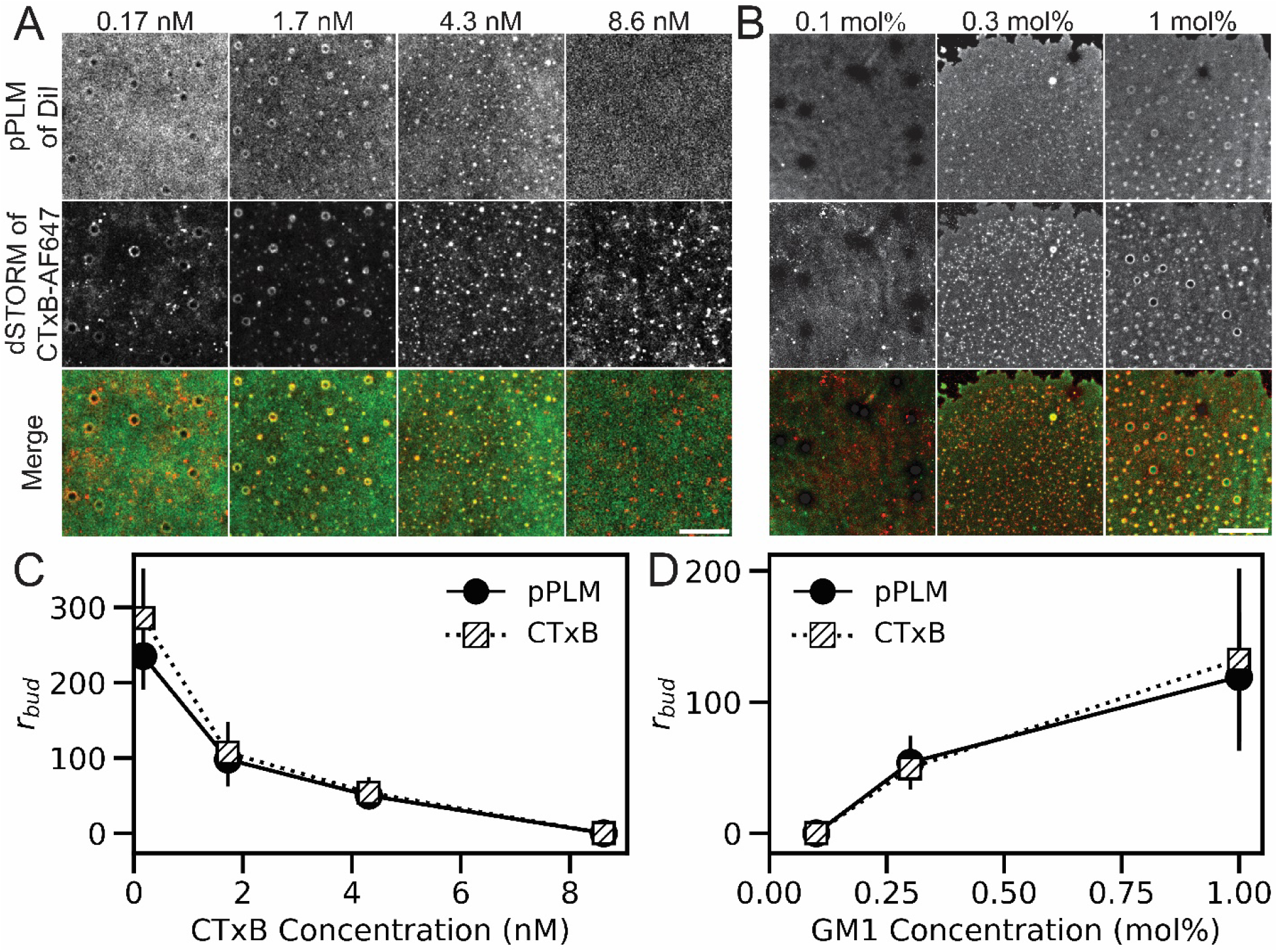
CTxB binding to supported bilayers containing GM1 induces buds that vary in size as a function of the concentration of both CTxB and GM1. (A) Planar supported lipid bilayers containing 0.3 mol% GM1 were incubated with CTxB concentrations ranging from 0.17 to 8.6 nM and subsequently imaged by pPLM and dSTORM. (B) Planar lipid bilayers containing varying GM1 concentrations were incubated with 4.3 nM of CTxB-AF647 prior to imaging using pPLM and dSTORM. The pPLM results quantify the membrane curvature, and dSTORM images show the CTxB distribution. Merged images of the DiI *(green)* and CTxB-AF647 *(red)* show strong co-localization of clustered CTxB at sites of membrane bud formation. Scale bars, 2 μm. (C) Measurements of membrane bud sizes from the super-resolution images of bilayers labeled with varying concentrations of CTxB reveal bud size decreases with increasing CTxB concentrations. (D) Bud size increases with increasing GM1 concentrations. Error bars represent the standard deviation of individual bud sizes from ≥ 3 repeats per condition.

Reconstructed images in the two channels revealed that regions of membrane budding were enriched in CTxB. Puncta of DiI localizations via pPLM indicated membrane buds and regions of the sample in which the membrane was perpendicular to the microscope coverslip (28). Interestingly, altering the GM1:CTxB ratio altered the radii of the membrane buds. Decreasing the GM1:CTxB ratio by increasing the concentration of CTxB from 0.17 to 4.3 nM, also decreased the average bud size from a maximum value of 285 ± 66 to 53 ± 20 nm as detected by dSTORM and 116 ± 68 to 49 ± 14 nm as reported by pPLM (Fig. 1). A similar result was observed when the GM1:CTxB ratio was varied by changing GM1 concentrations. In this case, the bud size increased with increasing concentrations of GM1. Measured values at 0.3 and 1 mol% were 50 ± 14 and 132 ± 70 nm via dSTORM and 53 ± 20 and 119 ± 50 nm via pPLM, respectively (Fig. 1). No membrane bending was detected at the lowest GM1-to-CTxB ratios examined, including either a high concentration of CTxB (8.6 nM) or a low concentration of GM1 (0.1 mol%). These findings reveal the GM1:CTxB ratio is an essential factor underlying the induction of membrane curvature, with high ratios giving rise to the formation of large buds and low ratios entirely failing to initiate membrane bending. Based on our results, we hypothesized that at low GM1:CTxB ratio, the predominant stoichiometry of membrane-bound CTxB is that of a single GM1 receptor bound to each CTxB, which does not induce membrane buds.

### Monovalent CTxB does not induce membrane budding

To test this hypothesis alternately, we determined the capacity of a monovalent variant of CTxB (mCTxB) to induce membrane curvature. Each mCTxB contains one native GM1-binding subunit and four subunits with G33D mutations that disrupt GM1 binding; each mCTxB can bind precisely one GM1 **(8)**. We examined the ability of 1.7 or 8.6 nM of mCTxB to induce budding in model membranes containing 0.3 mol% GM1. These concentrations of mCTxB were chosen to match the protein concentration and active GM1 binding site concentration, respectively, of the 1.7 nM wtCTxB condition that consistently induced membrane bending (Fig. 1). In contrast to the behavior of wtCTxB that contained 5 GM1 binding sites, no change in membrane topography was detected upon the addition of mCTxB to membranes at either concentration in model membranes (Fig. 2). To confirm the effect of valency on CTxB-induced curvature, we determined if mCTxB was capable of inducing curvature in intact giant unilamellar vesicles (GUVs). CTxB induced membrane invaginations on GUVs as expected **(3)**, whereas mCTxB caused no apparent change to the membrane shape (Fig. S1). Thus, the binding of CTxB to a single GM1 is not sufficient to bend membranes.

**Figure 2:**
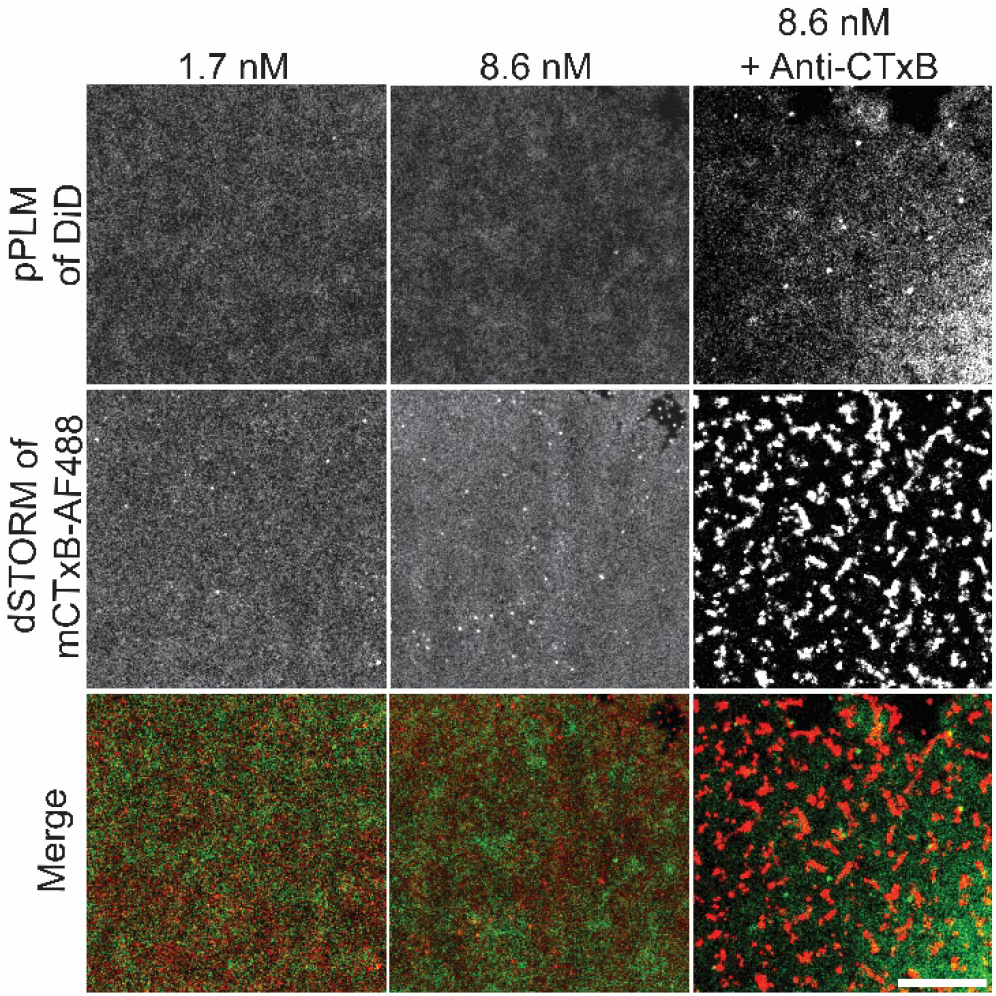
No membrane curvature is induced by the addition of mCTxB to GM1-containing membranes, even when it is crosslinked. Planar supported lipid bilayers containing 0.3 mol% GM1 were incubated with 1.7 nM mCTxB or 8.6 nM mCTxB. Cross-linking of the mCTxB was performed with a 1:100 dilution of anti-CTxB antibody. Membranes were subsequently imaged by pPLM and dSTORM. Crosslinking led to the formation of large-scale clusters in dSTORM images but nevertheless failed to generate significant levels of membrane budding. Lower concentrations of anti-CTxB demonstrate less mCTxB clustering and no membrane budding (Figs. S2 and S3). Merged images of the DiD *(green)* and mCTxB-AF488 (*red*) show no coincident puncta, indicating that mCTxB fails to form clusters or induce membrane curvature. Scale bar, 2 μm.

### Spatial organization of GM1 binding is necessary for inducing membrane curvature

We next tested the hypothesis that the spatial organization of binding sites on CTxB critically underlies the generation of membrane curvature. To test this idea, we asked if the clustering of monovalent CTxB by antibody cross-linking could override the requirement for multivalent binding by CTxB. For these experiments, membrane-bound mCTxB was crosslinked with varying amounts of anti-CTxB antibodies. The highest antibody concentrations used (1:100 dilution) induced clustering of mCTxB into punctate structures (Figs. 2, S2) and significantly slowed the diffusion of mCTxB (Fig. S3). However, no membrane bending was observed by pPLM at any anti-CTxB concentration examined.

We also tested if a general protein-induced clustering of lipids is sufficient to induce curvature. We labeled membranes containing varying concentrations of DPPE-biotin and crosslinked the lipids using streptavidin (Fig. S4). The tetravalent binding of streptavidin to DPPE-biotin yielded no apparent variations in the membrane topography as assessed with pPLM at any of the streptavidin or DPPE-biotin concentrations tested, consistent with previous reports (30). Thus, not all protein-induced lipid clustering is sufficient to bend membranes and the structural details of the GM1 binding sites in CTxB are essential for generating curvature.

### Single-particle tracking reveals complexes with high GM1-to-CTxB ratios diffuse extremely slowly

We used the diffusional behavior of CTxB-GM1 complexes to gain further insights into the stoichiometry of GM1 bound to CTxB in the membrane buds. To probe the dynamics of CTxB-GM1 complexes, the single-molecule blinking data used in the reconstruction of superresolution images for wtCTxB and mCTxB were analyzed using high-throughput single-particle tracking. To eliminate the possibility of topographical effects slowing diffusion, we excluded any single-particle trajectories that were coincident with punctate localizations in pPLM. As a control, an identical analysis was performed for DiI. Minimal effects of CTxB binding were observed on DiI diffusion. Under all experimental conditions tested, DiI exhibited diffusion consistent with a single population diffusion coefficient *(D)* of 0.46 ± 0.2 μm^2^/sec.

mCTxB, like DiI, demonstrated a single population of Brownian diffusers with a characteristic *D* of 0.74 ± 0.54 μm^2^/s. In contrast, wtCTxB displayed heterogeneous, multipopulation distributions of diffusion coefficients. Greater GM1 availability increased the abundance, but not the speed, of the slower moving populations of wtCTxB (Figs. 3, S5).

**Figure 3:**
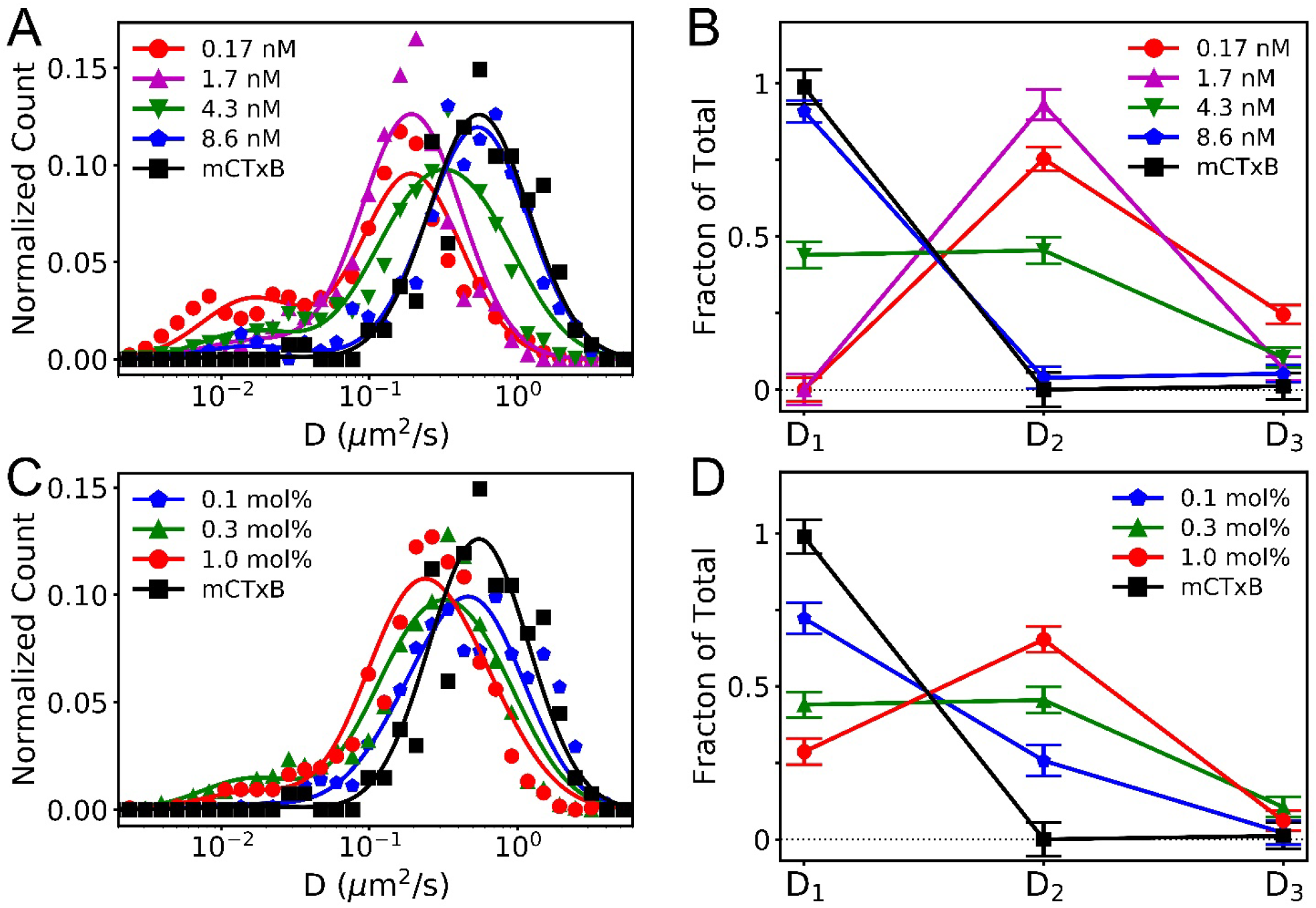
Single-particle tracking of CTxB reveals a distribution of diffusion rates suggestive of the presence of three distinct subpopulations of CTxB bound to differing numbers of GM1s. Single-particle tracking analysis was conducted on individual, membrane-bound CTxB molecules in bilayers containing constant 0.3 mol% GM1 and varying CTxB concentrations (A, B) or constant 4.3 nM CTxB and varying GM1 concentrations (C, D), as also imaged in Fig. 1. For comparison, single-particle tracking was also performed for 1.7 nM mCTxB. Histograms report the distribution of *D* values obtained under each condition (A, C). The histograms were fit using Eq. 1 to quantify the diffusion coefficient and frequency of the three observed CTxB subpopulations (B, D). Fits of individual subpopulations are shown in Fig. S5. The slower diffusing subpopulations were more abundant at lower CTxB concentrations or higher GM1 concentrations, which correspond to higher GM1:CTxB ratios and to greater membrane bending.

To discern the underlying subpopulations, the experimentally acquired distributions of *D* were fit to a probability distribution (*P*) composed from a sum of subpopulations of diffusers (Eq. 1) with each subpopulation contributing a log-normal distribution of *D* (Eq. 2).

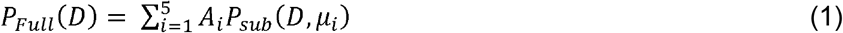

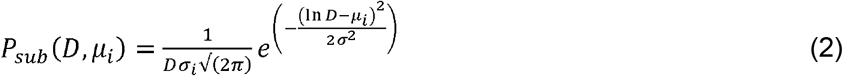

The width of the log-normal distribution (*σ*) was assumed to be as theoretically expected from the distribution of trajectory lengths analyzed and constant for all subpopulations (31); *σ* = 0.78. The mean diffusion coefficient for each subpopulation *(D_i_)* is equal to exp(*μ_i_*+*σ*^2^/2).

Three subpopulations were sufficient to fit all our data of CTxB diffusing on model membranes. *D_i_* values of *D*_1_ = 0.75, *D*_2_ = 0.26, and *D*_3_ = 0.022 μm^2^/s were identified by fitting the eight experimentally acquired data sets simultaneously (Fig. S5). We interpret *D*_1_ as the mean diffusion coefficient for CTxB with *i* GM1 per CTxB, *i.e., D*_1_, *D*_2_, and *D*_3_, corresponding to the diffusion rates of CTxB bound to 1,2, and 3 GM1, respectively. The subpopulations of 4 or 5 GM1-to-CTxB were not apparent in our super-resolution observation (*i.e., A*_4_ = *A*_5_ = 0). The sum of the three fit *A_i_* values were normalized to represent the relative fraction of CTxB with 1,2, or 3 GM1 per CTxB in each experiment (Fig. 3B, D). As expected, the experiments with mCTxB, concentrated CTxB, or dilute GM1 each yielded *A*_1_ close to unity, confirming one GM1 per CTxB. In contrast, with decreasing CTxB concentration or increasing GM1 concentration, the subpopulations that represented more GM1-per-CTxB became prevalent.

In addition to confirming the presence of multiple stoichiometries of CTxB-GM1 complexes, the results of these experiments show the diffusion coefficients do not scale linearly with the number of bound GM1 molecules. The dramatic slowing of CTxB diffusion with an increasing GM1:CTxB ratio suggests that CTxB either progressively penetrates the membrane or induces local membrane deformations that slow their diffusion. Taking these results together with the PLM results, we conclude that the binding of CTxB to just two GM1 was sufficient to induce membrane curvature. Further increasing the binding stoichiometry resulted in increasing the radii of the buds.

### Varying ceramide structure or cholesterol content had minimal effects

The ceramide structure of GM1 functions as a key regulator of intracellular trafficking of both GM1 itself as well as CTxB-GM1 complexes (32), suggesting it plays an essential function in sorting as the toxin moves from one intracellular compartment to another. Furthermore, GM1 and Gb3 structure has been directly linked to curvature induction by SV40 and STx, respectively (3–5). We thus tested the hypothesis that the fatty acyl chains of the ceramide domain of GM1 are important for toxin-induced membrane curvature. Three custom gangliosides were used to address this question, recognizing that longer and saturated acyl tails are more correlated with ordered lipid phases (Fig. S6). In our model systems, membrane bending was detected at similar levels for GM1_16:1_, GM1_18:1_, or GM1_18:0_ upon the addition of CTxB, and there was no systematic change in the radii of the buds correlated with the expected lipid phase preferences of the tails (Fig. 4).

**Figure 4:**
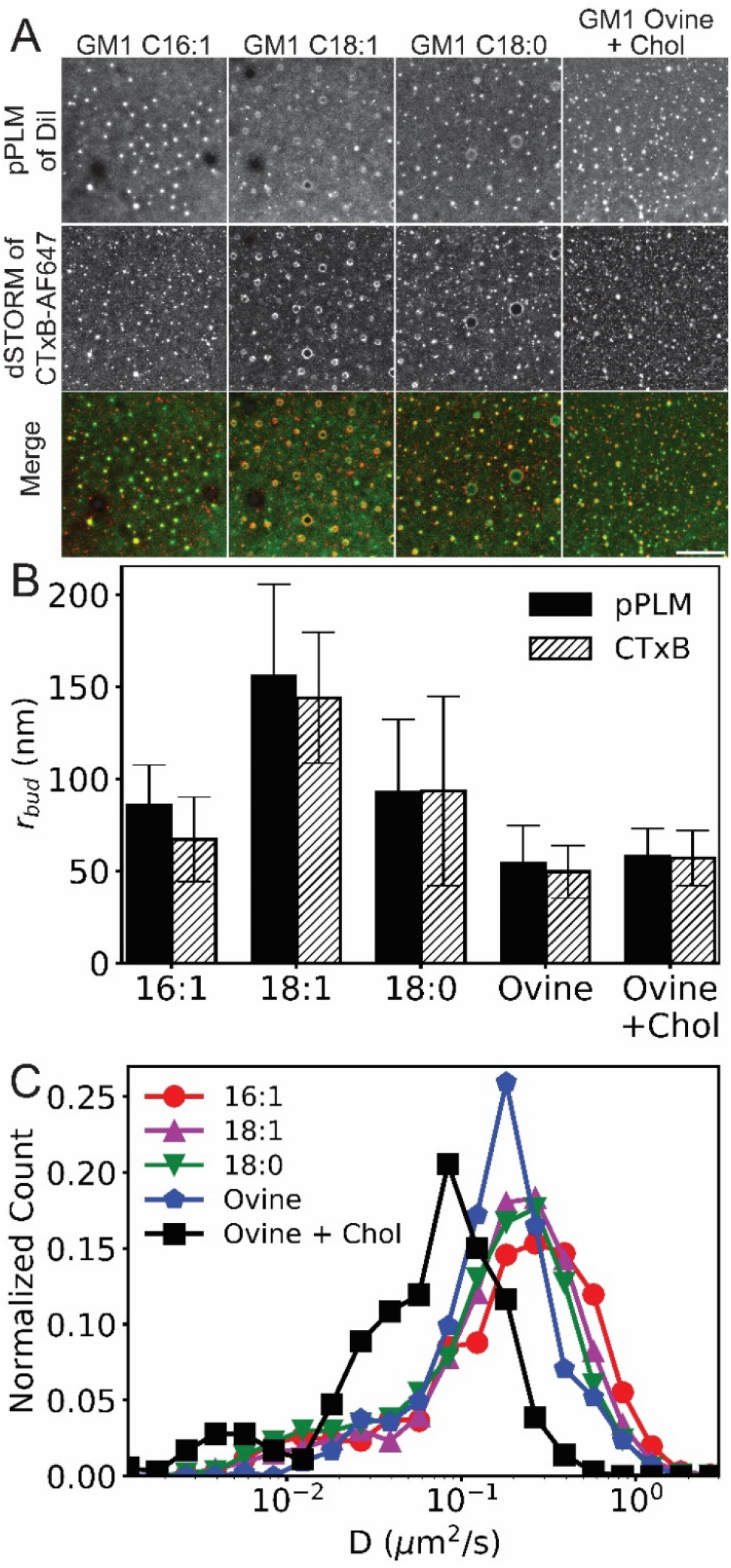
Curvature induction in response to CTxB binding is not dependent on the GM1 acyl chain structure or affected by the presence of cholesterol. 4.3 nM CTxB was bound to planar lipid bilayers containing 0.3 mol% of synthetic GM1 with acyl chains of 16:1, 18:1, or 18:0 or commercially available ovine GM1 plus 30 mol% cholesterol. (A) Samples were subsequently imaged using pPLM to detect membrane curvature or dSTORM to image CTxB clustering. Merged images of the DiI (*green*) and CTxB (*red*) show strong co-localization of clustered CTxB at sites of membrane bud formation regardless of the GM1 acyl tails or cholesterol content. Additional CTxB concentrations are shown for cholesterol-containing membranes in Fig. S7. Scale bar, 2 μm. (B) Curvature was quantified by measuring the mean and standard deviation of membrane bud sizes from three repeats. The bud sizes for GM1_18:1_ were significantly larger than that of GM1_Ovine_ in the presence or absence of cholesterol (*p*<0.05), but no other statistical differences were present. (C) Histograms of the distribution of *D* from single-molecule trajectories of CTxB bound to planar membranes containing the indicated GM1 species or a combination of ovine GM1 and 30 mol% cholesterol. The CTxB diffusion slowed with increasing length and saturation of the GM1 acyl tails or in the presence of cholesterol.

Cholesterol levels strongly regulate endocytosis of CTxB (33). We thus wondered if cholesterol levels might control the ability of CTxB to induce membrane curvature. To test this, we added 30 mol% cholesterol to the supported bilayers. The bud radii obtained in samples with and without cholesterol were 50 ± 14 and 57 ± 15 nm via dSTORM and 54 ± 14 and 58 ± 15 nm via pPLM, respectively, when 1.7 nM CTxB and 0.3 mol% GM1_Ovine_ were used (Fig. 4). Just as in the absence of cholesterol, in the presence of cholesterol, higher CTxB concentrations resulted in less membrane bending (Fig. S7). Thus, neither the ceramide structure nor cholesterol inclusion impacted the capacity of CTxB to induce curvature in artificial membranes under the conditions of our experiments.

Diffusion coefficient of CTxB when bound to GM1_16:1_, GM1_18:1_, GM1_18:0_, GM1_Ovine_, or in the presence of cholesterol was 0.29 ± 0.27, 0.25 ± 0.23, 0.22 ± 0.21, 0.21 ± 0.17, and 0.081 ± 0.08 μm^2^/sec, respectively. The trend observed in the distribution of diffusion coefficients is consistent with slower diffusion for longer or more saturated acyl tails in addition to by increasing acyl tail ordering through the addition of cholesterol (Fig. 4C).

### GM1 binding stoichiometry affects membrane bending in cells

Finally, we tested if the requirement for multivalent binding by CTxB to induce curvature is also conserved in cellular membranes. We assayed for curvature induction upon CTxB binding in COS-7 cells using polarized localization microscopy (28). 1.7 nM of CTxB or mCTxB was applied to live COS-7 cells stained with DiI (Fig. 5). Cells labeled with CTxB displayed punctate accumulations of DiI and CTxB, with a mean bud radius of 220 ± 160 nm and 260 ± 150 nm as detected by pPLM and dSTORM, respectively. There were no apparent puncta in cells labeled with both DiD and mCTxB, indicating that multivalent GM1 binding was also required to generate membrane curvature in cells.

**Figure 5:**
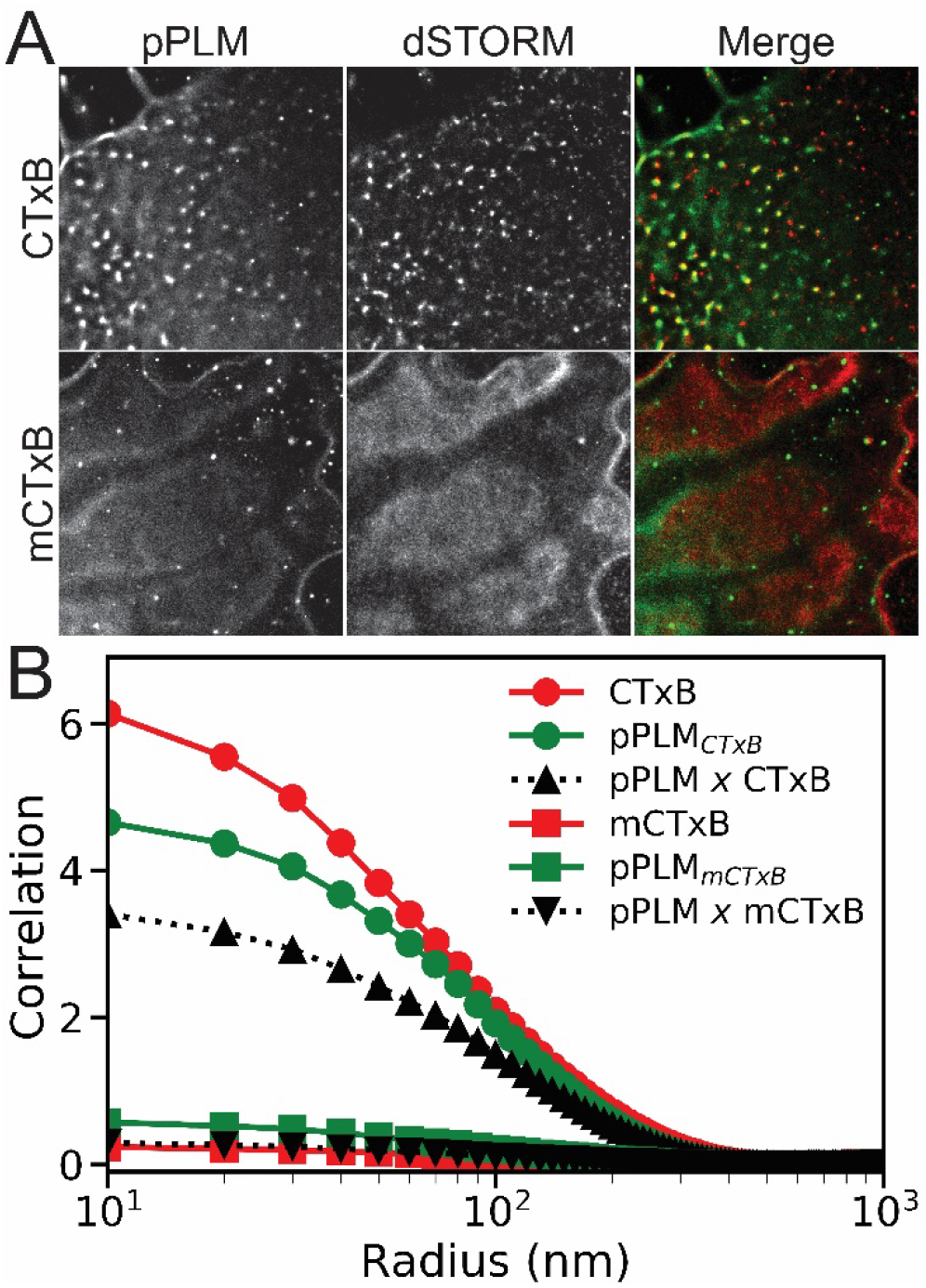
CTxB clusters and induces curvature in the plasma membranes of living cells, whereas mCTxB undergoes minimal clustering and fails to generate curvature in cell membranes. COS-7 cells were labeled with DiI or DiD and 1.7 nM of CTxB or mCTxB. They were subsequently imaged using pPLM and dSTORM. Merged images show areas where clustered CTxB or mCTxB (*red*) colocalize with sites of induced curvature (*green*). Scale bar, 2 μm. (B) Spatial autocorrelations were performed to quantify the magnitude of the clustering of CTxB (*red circles)* or mCTxB (*red squares)* and the magnitude of nanoscale membrane budding induced by either CTxB *(green circles)* or mCTxB *(green squares).* Cross-correlation analysis was performed to quantify the magnitude of the sorting of CTxB *(black triangles)* or mCTxB *(black inverted triangles)* to sites of membrane bending. Cells labeled with CTxB show an increase in nanoscale membrane bending that is spatially correlated to the CTxB clustering, whereas those labeled with mCTxB show minimal membrane curvature induction and negligible sorting of the mCTxB to curvature sites.

To quantify this difference, we performed spatial cross-correlation analysis and assessed the characteristic abundance, size, and colocalization of the punctate structures (Fig. 2). Significant cross-correlation between pPLM and CTxB was observed, demonstrating that the protein localization was spatially coincident with curved membranes with a characteristic radius of 85 to 105 nm. In contrast, pPLM and mCTxB showed negligible cross-correlation. The amplitude of spatial autocorrelations of CTxB and pPLM after CTxB addition were 14-fold higher than those for mCTxB and pPLM after mCTxB addition. We conclude there is a substantial increase in nanoscopic membrane bending events when CTxB, but not mCTxB, is added to cells (Fig. 5B).

To confirm the differential membrane binding induced by CTxB and mCTxB binding to cells, single-molecule diffusion coefficients were again examined (Fig. S8). On live cells, CTxB and mCTxB displayed *D* of 0.33 ± 0.40 and 0.17 ± 0.20 μm^2^/s, respectively.

## DISCUSSION

The clustering of lipids by AB_5_ toxins and SV40 virus is thought to induce compression of the lipids as a means of bending membranes (3, 4, 34) via a process linked to the geometry of receptor binding sites (16, 35). However, much of this process warrants further characterization, including the role of the clustering and leaflet asymmetry in the generation of curvature (36). In this study, we investigated the factors that drive membrane bending by CTxB, a prototypical cargo for clathrin-independent endocytosis, using super-resolution polarized localization microscopy.

There are two important considerations that must be emphasized to understand our results. First, this curvature is induced on supported lipid bilayers (SLBs), which have adhesion to the underlying support that reduces the diffusion of lipid domains, imposes membrane tension, and reduces shape fluctuations (37). Our method of creating SLB patches via GUV fusion facilitates membrane bending by apparently trapping the center of the patch with a lower tension and less adhesion than the perimeter of the patch *(i.e.,* Fig. 1B) (17). Secondly, CTxB is exposed to the distal side of the SLB such that membrane curvature can only form by bending the membrane away from the substrate and towards the CTxB. This direction of bending is the opposite of that observed on cells and GUVs *(i.e.,* Fig. S1) but potentially consistent with the preference for CTxB to bind to negative principal curvatures (Fig. S9) (17). The SLBs used in this study provide control of the lipid and protein constituents such that the mechanisms and stoichiometric dependence CTxB membrane bending could be ascertained.

Our data support the hypothesis that toxin-induced glycolipid crosslinking induces membrane curvature in both model membranes and cellular environments. We also show that the multivalent binding of GM1 by CTxB is necessary for the membrane curvature and experimentally demonstrate the importance of GM1-to-CTxB stoichiometry in the membrane bending capability of CTxB. Importantly, lipid scaffolding on its own is not sufficient to induce membrane bending and more dramatic membrane shape changes occurred at lower CTxB concentrations. Thus, the mechanism by which CTxB induces curvature likely does not arise from protein crowding effects (38) but does arise as a consequence of the spatial organization of glycolipids induced by multivalent binding to pentavalent CTxB.

We also determined the likely stoichiometry of GM1 binding to CTxB by fitting the histogram of the diffusion rate to a sum of log-normal Gaussians (Eq. 1). This single-particle diffusion analysis directly reports on the number of GM1s bound to CTxB and enables correlation of membrane bending activity to structured lipid crosslinking. While mCTxB engineered with a single binding site had only a single population of diffusers, wtCTxB had multiple species at the concentrations tested. Increasing the wtCTxB concentration resulted in faster wtCTxB diffusion, allowing us to exclude CTxB aggregation as a likely mechanism underlying our observations.

The distribution of *D* of wtCTxB was well fit with three populations representing CTxB bound to 1,2, or 3 GM1 per CTxB (Figs. 3, S5). This interpretation is consistent with computational simulations (39), plasmonic nanocube membrane binding assays (40), flow cytometry (41), and mass spectrometry (42) that show concentration-dependent binding stoichiometries and that saturating the CTxB binding sites is unexpected for the concentrations used here. The CTxB concentration-dependent dissociation rate constants measured by flow cytometry show that 20 nM of CTxB results in 1 GM1 per CTxB with an off rate of 5.2 min and lower concentrations of CTxB displayed slower off rates (41). A single CTxB maintains a binding conformation for a longer duration than our single-molecule trajectories. Maximum membrane coverage occurred when ≤1 mol% GM1 was exposed to 20 nM CTxB (40, 41) and each GM1 was bound to a separate CTxB. Prior studies are consistent with our observed variations in GM1:CTxB ratios.

The diffusion rates observed here are consistent with the variety of previously published average diffusion rates for CTxB (17, 20, 43–49). The diffusion of mCTxB and the fastest CTxB diffusion are consistent with the diffusion of unbound GM1 (50), implying that monovalent CTxB binding provides no significant membrane disruption *(i.e.,* curvature) or resistance to GM1 diffusion. A non-linear slowing of the diffusion coefficient was observed upon increasing the stoichiometry of GM1 per CTxB. CTxB bound to one GM1 diffuses 34x faster than CTxB bound to three GM1. This slowdown with lipid binding is more than is expected by the free draining limit (51, 52) and reveals that increased GM1 per CTxB is correlated with membrane perturbations in addition to GM1 cross-linking (53). Slowed CTxB diffusion reduces the probability of saturating the GM1 binding sites because protein and glycolipid diffusion affects their association rate, which is key in determining the observed binding stoichiometries for SV40 (9) and CTxB (54).

Based on these findings, we propose a model for how the stoichiometry of GM1 binding to cholera toxin can be mechanistically linked to curvature generation (Figs. 5, S9). With zero GM1 bound per CTxB, the CTxB is not membrane-bound, and there is no curvature generation. With one GM1 bound, the CTxB is attached to the membrane, but there is no curvature generation (Fig. 6A). This single-bound-GM1 conformation likely includes the CTxB sitting tilted on the membrane to juxtapose the single GM1 binding pocket close to the lipids with minimal membrane shape dependence, as observed previously in computational models (16, 39, 55, 56). With two GM1s bound, however, CTxB has a strong preference for negative curvature along one dimension and a weak preference for negative membrane curvature along the perpendicular dimension (Fig. 6B). Further increasing the GM1:CTxB ratio increasingly perturbs the membrane, giving rise to an enhanced preference towards negative curvature, which is more available on larger membrane buds (Fig. S9).

**Figure 6:**
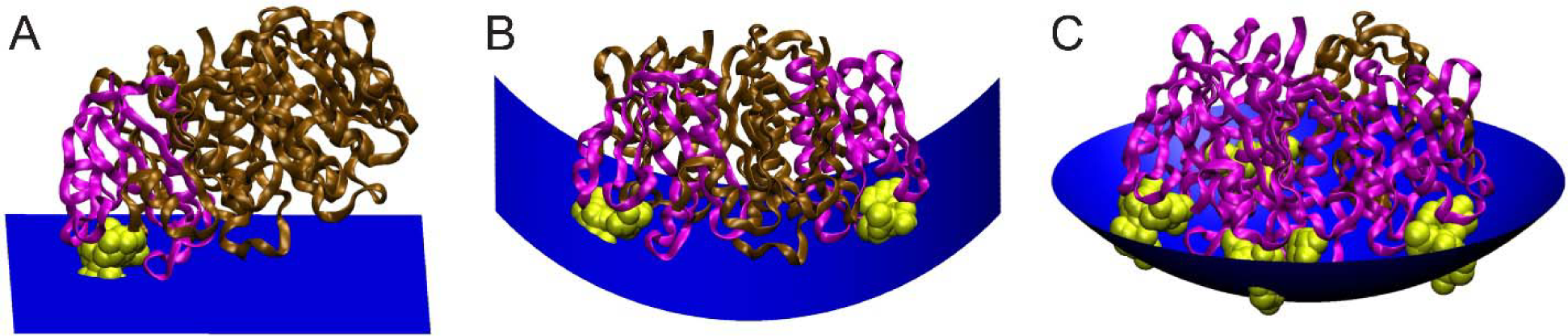
Model for how the stoichiometry of binding of CTxB to GM1 generates membrane curvature. The ratio of CTxB subunits bound (*pink*) and unbound *(brown)* to GM1 *(yellow)* affects the shape of the membrane surface *(blue).* (A) Binding CTxB to a single GM1 has no effect on membrane curvature. (B) Binding CTxB to two GM1s generates negative curvature in one dimension, such as occurs at the neck of a membrane bud (Fig. S9). (C) Binding of CTxB to 3 or more GM1s induces negative membrane curvature in two dimensions. This results in a wrapping of the membrane around the CTxB, such as within an endocytic pit. The schematics were created in VMD (75) by building upon a crystal structure of CTxB (76).

Our results provide insights into the steps in the endocytosis and trafficking of cholera toxin that rely on the ability of the toxin to induce curvature. CTxB can be internalized through different endocytic pathways (21, 25, 57), including via tubular clathrin-independent carriers generated in response to toxin binding (3, 4, 58–60). Our results support the notion that wild type toxin is inherently capable of initiating membrane bending events and that curvature induction requires the toxin to bind multiple GM1. However, monovalent variants of CTxB are efficiently sorted into surface-attached endocytic tubules (58). Furthermore, “chimeric” toxins with differential number of GM1 binding sites can undergo endocytosis albeit less efficiently compared to wild type cholera toxin (61), and binding of CTxB to a single GM1 is sufficient to complete the intoxification pathway (8). Given that the ability of CTxB to induce curvature is dependent on multivalent binding to GM1, these findings suggest the CTxB-induced curvature is not required for sorting of toxin into nascent endocytic structures, entry of the toxin into cells by endocytosis, or directing its intracellular transport into pathways that leads to toxicity. However, the conditions in which CTxB induces curvature to enhance the efficacy of the toxin, perhaps due to the recruitment of ancillary host proteins that facilitate endocytosis (59, 60) or the increased affinity of the CTB-GM1 complex for curved membrane structures (*i.e.*, sorting tubules or endocytic pits).

It is well known that the ceramide structure is important for intracellular lipid sorting pathways and influences the ability of AB_5_ toxins to bend membranes. Unsaturated GM1 is a better receptor for retrograde trafficking of cholera toxin to the endoplasmic reticulum than saturated GM1 (32). In the case of another AB_5_ toxin, Shiga toxin, the side chains of its glycolipid receptor are required to be unsaturated for tubulation in artificial vesicles (4). Furthermore, long-chain but not short-chain GM1 is essential for SV40 to generate membrane invaginations (3). Motivated by these findings, we tested the hypothesis that the ceramide structure of GM1 would be relevant to the initial membrane bending process using both synthesized and naturally occurring GM1 variants. Our experiments showed there was minimal change in membrane topography in model membranes containing GM1 variants. This result suggests that CTxB-induced membrane bending does not rely on the acyl chain identity of GM1, at least within the range of the ceramide structures tested in this study. Our experiments were performed only in artificial membranes, and we do not rule out the possibility that ceramide structure may impact the ability of CTxB to generate membrane curvature in the cell surface or intracellular membranes of intact cells. In phase-separated vesicles, for instance, ceramide tail structure is critical for raft partitioning of both uncrosslinked and CTxB-crosslinked GM1 (32). Thus, ceramide structure could impact the raft association of cross-linked GM1, which is credited with the membrane bending and internalization of CTxB in cells (62–65). Variation in the ceramide tail structure may alter the composition and acyl-tail order of GM1-rich nanoscopic domains (43, 66, 67). Accordingly, the spontaneous recruitment of liquid-order preferring lipids and the line tension surrounding the GM1-rich domain are likely to depend on the ceramide structure. Given the importance of ceramide structure in various physiological contexts, we hypothesize that the role of the GM1 ceramide domain is to recruit the downstream endocytic machinery, such as clathrin, caveolin, flotillin, or dynamin.

These results also have implications for the use of cholera toxin as a marker for the liquid-ordered (raft) lipid phase. CTxB is not a non-perturbing marker for ordered domains; it can induce phase separation in membranes close to a demixing point (7, 63), stabilize ordered phases (7, 68), sort to already curved membranes (13, 17, 18), and induce membrane curvature (3, 17). Our current findings reveal that any measurements of the impact of CTxB on a membrane require precise control of the CTxB and GM1 concentrations because the diffusive, sorting, and curvature-inductive properties of CTxB depend on its binding stoichiometry.

## MATERIALS AND METHODS

### Custom GM1 synthesis

GM1 variants in the ceramide were synthesized in DMSO solvent containing 1 μmole lyso-GM1, 10 μmoles fatty acid, 10 μmoles of dicyclohexyl-carbodiimid and 10 μmoles of sulfo-N-hydroxysuccinimide for 3 hours at room temperature, as described previously (32). The DMSO was evaporated using a Speed Vac (Thermo Fisher Scientific) and purified following established methods via a silica column, preparative-TLC plates, and reversed-phase Bond-Elut cartridges (69). The products were analyzed by electrospray ionization mass spectrometry with a quadrupole orthogonal time-of-flight instrument (Q-o-TOF MS, Q-Star Pulsar *i,* Applied Biosystems, Toronto, Canada).

### SLB preparation and formation

SLBs were made from bursting GUVs that included 1-palmitoyl-2-oleoyl-sn-glycero-3-phosphoch (POPC; Avanti Polar Lipids), 1,1’-dioctadecyl-3,3,3’,3’-tetramethylindocarbocyanine perchlorate (DiI; Life Technologies, Carlsbad, CA), and GM1. Commercially available Ovine GM1 (Avanti Polar Lipids) or GM1 with custom acyl tails were used. All SLBs were primarily POPC with cholesterol (Sigma-Aldrich) only included when indicated. All SLBs included 0.3 mol% of DiI or DiD for CTxB-AF647 or mCTxB-AF488 experiments, respectively. For streptavidin-biotin binding experiments, GM1 was replaced with DPPE-Biotin (Avanti Polar Lipids).

GUVs were prepared by electroformation, as described previously (70). Briefly, a hydration solution of 200 mM sucrose was added to the dried lipids on an ITO slide. A chamber was created with a silicone frame and a second ITO plate enclosing the lipids and sucrose solution. The two ITO plates were connected to a sine wave function generator at 2 V_peak-to-peak_ and 10 Hz for 3 hr at 50 °C.

### wtCTxB and mCTxB

Alexa Fluor 647-wtCTxB (CTxB) was obtained from Invitrogen. Alexa Fluor 488-mCTxB was made as described previously (8, 62). Briefly, an *E. coli* expression strain containing the plasmids encoding native CTxB-G33D (non-binding mutant) and C-terminal GS-H6-tagged wtCTxB was induced. Each subunit was secreted to the periplasm where mixed pentamers assembled. This mixture of assembled was purified from a cell extract by Talon affinity chromatography and separated into individual species by ion-exchange chromatography. Further purification was performed by cationic resin, HQ20 anion exchange column, and anion exchange prior to buffer exchange into PBS by ultrafiltration using an Amicon Ultra-4 (Millipore) 10 K cutoff centrifugal filter. Labeling was performed by reacting Alexa Fluor 488-succinimidyl ester (Invitrogen) to 300 μg mCTxB in 100 mM sodium bicarbonate buffer pH 8.3 for 1 h under stirring at room temperature and purified using provided size exclusion chromatography resin. DiI replaced DiD in the experiments on model membranes with the mCTxB-AF488 addition to decrease channel crosstalk. Model membranes were labeled by CTxB and mCTxB with a 0.5 min of incubation, after which the unbound CTxB or mCTxB was rinsed away. The resulting membrane budding showed no change 20 min after CTxB was added; imaging was performed approximately 60 min after CTxB addition.

### Cell Labeling

DiI-C18 or DiD-(Sigma-Aldrich) was dissolved in ethanol at 1 mM. Cells were rinsed twice with HEPES-based cell imaging buffer (Sigma-Aldrich), and the 1 mM DiI-C18 stock solution was added to COS-7 cells in HEPES at 1:100 dilution for 10 sec prior to washing with excess HEPES buffer. Cells were exposed to 1.7 nM CTxB-AF647 or mCTxB-AF488 for 5 min at room temperature prior to rinsing twice with HEPES imaging buffer prior. 7 individual cells were analyzed per condition.

### Anti-CTxB antibody addition

mCTxB crosslinking was induced by the addition of anti-CTxB antibodies (Calbiochem, 227040 goat pAb). Antibodies were added to the membrane at the 90 min time point following mCTxB addition at final dilutions of 1:10,000, 1:1000, 1:500, and 1:100 and allowed to incubate for 30 min prior to imaging.

### Imaging optics

Dual-color imaging was performed with a custom IX83 (Olympus). The excitation light polarization was controlled by a liquid crystal wave plate (LCC1111-A; Thorlabs). The excitation lasers were steered through a periscope and a 4-band notch filter (ZET405/488/561/640m; Chroma) onto the back of a 100x, 1.49 NA objective (Olympus). The emitted light passes through emission filters (BrightLine single-band bandpass filters; Semrock) within an OptoSplit with a 2.5X magnification lens (Cairn Research) before being collected by an iXon-897 Ultra EMCCD camera (Andor Technology).

Samples were exposed to high laser power (>80 mW) with λ_ex_ = 488 nm (mCTxB-AF488), λ_ex_ = 561 nm (DiI), or λ_ex_ = 647 nm (CTxB-AF647 or DiD) to provide a steady rate of photo-switching by the fluorescent probes. Sample imaging was performed 90 min after CTxB addition. 40,000 frames of blinking fluorophores were acquired for each two-color channel with an 18 ms exposure time at a 50 Hz frame rate.

### Imaging buffer

PLM and dSTORM were performed on SLBs surrounded by an oxygen-scavenging buffer (150 mM NaCl, 50 mM TRIS, 0.5 mg/mL glucose oxidase, 20 mg/mL glucose, 40 μg/mL catalase, and 1% β-mercaptoethanol at pH 8). This buffer encourages the reversible blinking of fluorophores in a low oxygen concentration. Buffer proteins were purchased from Sigma-Aldrich (St. Louis, MO), and salts were purchased from Thermo Fisher Scientific.

### Single-molecule localization microscopy

Single-fluorophore localizations were identified by the ImageJ plugin ThunderSTORM using bilinear interpolation to achieve sub-pixel precision (71). Single-fluorophore localizations with intensities <100 photons or location uncertainty >45 nm were excluded from the analysis. The separate channel images were overlaid via a custom-made MATLAB routine (The MathWorks, Natick, MA) by aligning the TetraSpeck nanoparticles (100 nm diameter; Life Technologies) that were visible in all channels and maximizing the image cross-correlation. The bright TetraSpecks localizations reduced the probability of finding a single fluorophore in their immediate proximity. The bright TetraSpeck localizations were hidden from the reconstructed images for clarity and removed prior to analysis of the single-molecule data.

### Bud identification and size evaluation

The detection of membrane buds in each color channel was performed by a custom-made MATLAB program that masks regions with a low local density of localizations to identify regions of curvature as described previously (28). 2D Gaussian fitting for small buds (<150 nm) and 2D toroidal fitting for buds (>150 nm) that had a ring shape were performed for center estimation. The size of the bud was calculated by evaluating the mean distance of all localizations at the bud location while considering the additional localizations from the surrounding flat membrane, as done previously (28). The number of localizations expected within a radius R if no bud was present (*N_SLB_*); *N_SLB_*= *ρπR*^2^ = *N_all_-N_bud_*, where *ρ* is the uniform density of localizations on a flat membrane, *N_bud_* is the number of extra localizations due to the presence of the bud. The mean distance of the localizations from the center of the bud expected for the flat SLB within *R* is 2*R*/3. By analyzing all collected localizations within R and subtracting the expected localizations from the flat SLB, the size of a single budding event (*r_bud_*) was calculated according

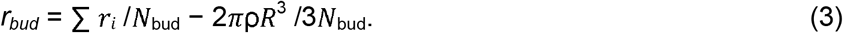

### Single-particle tracking

Single fluorophores were analyzed to reveal the diffusion rates of the lipid and toxin. Blinking fluorophores were linked based on their proximal distance in consecutive frames via u-track (72), and single trajectories that lasted ≥15 frames were further analyzed. The slope of the single-molecule MSD vs. Δ*t* for Δ*t* from 2 to 4 frames were fit to obtain the single-molecule diffusion rates according to

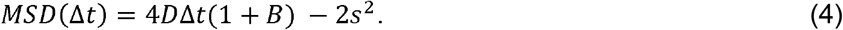

The localization imprecision (*s*) and the motion blur coefficient (*B*) were equal to 15 nm and 1/6, respectively, for the imaging parameters used here (73).

### Image correlation analysis

Spatial cross-correlation of two images, *I_1_* and *I_2_*, were computationally calculated according to Eq. 5 and azimuthally averaged, as done previously (74).

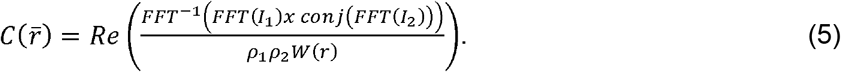

FFT represents the fast Fourier transform, *conj* represents the complex conjugate, and *Re* represents taking only the real component. The correlations are normalized by the average localization density (*ρ*) and the azimuthally averaged autocorrelation of the region of interest (*W*). The autocorrelation of an image, *I_1_*, was calculated from Eq. 5 by setting *I_2_* = *I_1_*.

### Data availability

The data that support the findings of this study are available from the corresponding author upon reasonable request.

## Author Contributions

A.M.K., K.R., W.I.L., A.K.K., and C.V.K. designed the research. A.M.K. performed the imaging. A.M.K. and C.V.K. performed the data analysis. A.M.K., K.R., W.I.L., A.K.K., and C.V.K. wrote the paper.

## ACKNOWLEDGMENTS

We thank Eric Stimpson, KeVaughna Patrick, Sonali Gandhi, Meshari Al Rafidi, Xin Xin Woodward, and Stephanie Schmieder for helpful discussions and comments. AMK was funded by a Thomas C. Rumble Fellowship Award. Financial support was provided by Wayne State University laboratory start-up funds and the Richard J. Barber Interdisciplinary Research Program. KR was supported in part through by Children’s Hospital of Pittsburgh of the UPMC Health System. This material is based upon work supported by the National Science Foundation under grant no. DMR-1652316 and the National Institute of General Medical Sciences of the National Institutes of Health under Award Number R01GM106720. The content is solely the responsibility of the authors and does not necessarily represent the official views of the National Institutes of Health.

## SUPPLEMENTAL FIGURES

**Figure S1:**
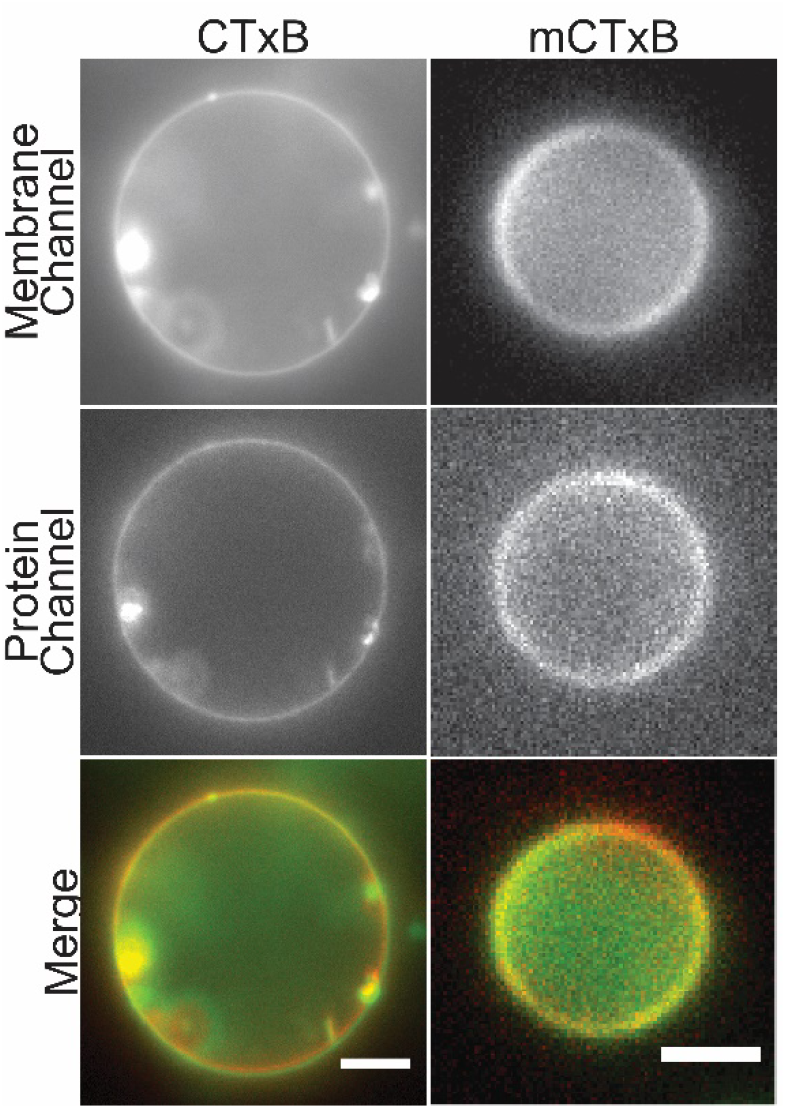
Binding of CTxB, but not mCTxB, to GUVs induces the formation of tubular invaginations. GUVs composed of POPC and 0.3 mol% GM1 were labeled with 0.3 mol% DiI or DiD and incubated for 20 min in the presence of either 1.7nM CTxB or 1.7nM mCTxB prior to imaging with wide-field fluorescence microscopy. Scale bars, 5 μm.

**Figure S2:**
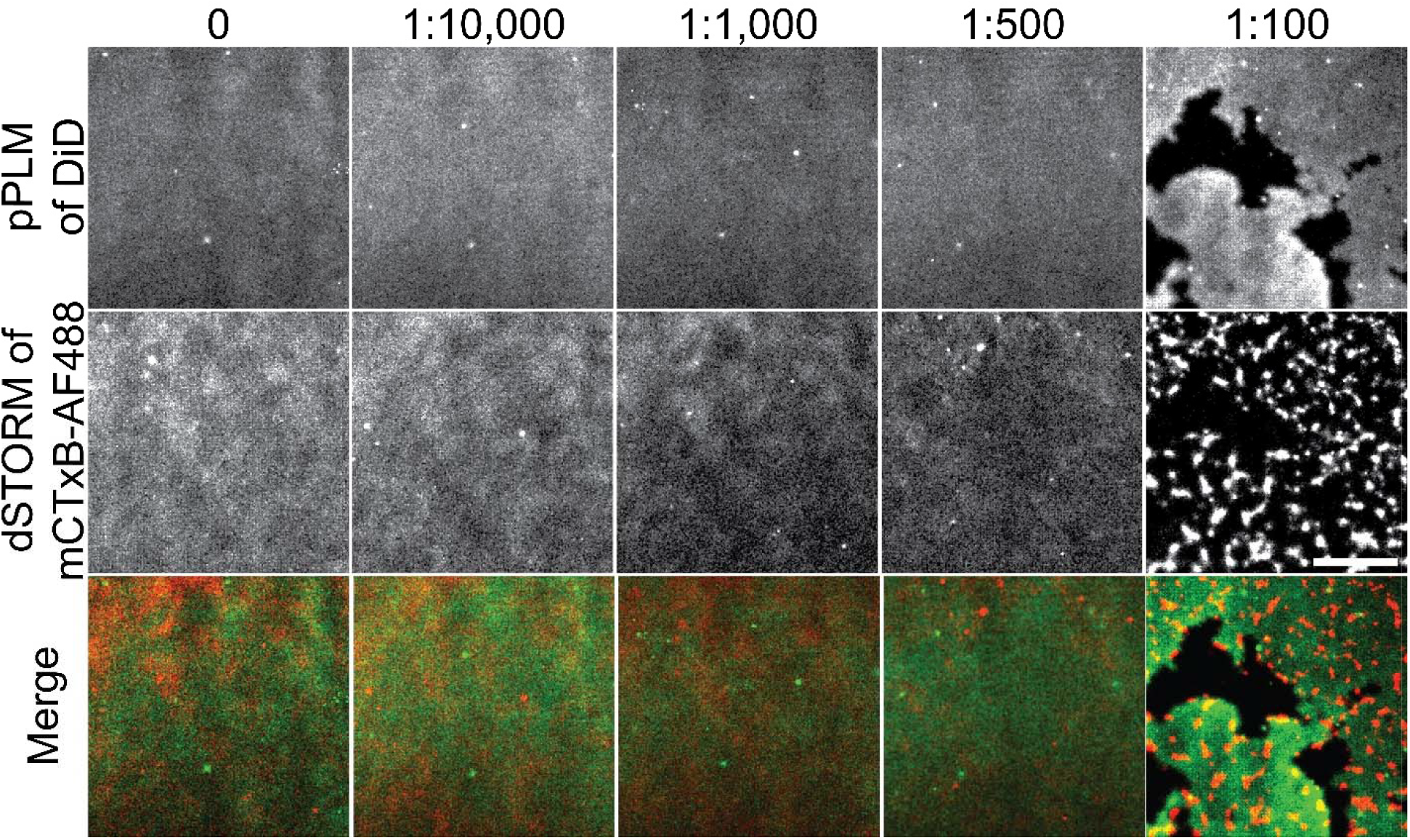
Antibody-induced crosslinking of mCTxB fails to induce membrane budding. Planar lipid bilayers containing 0.3 mol% GM1 were labeled with 8.6 nM mCTxB followed by either buffer alone or buffer containing the indicated dilutions of anti-CTxB antibody. They were subsequently imaged by pPLM (A-E) and dSTORM (D-F). Note that the clustering of the mCTxB was readily observed at an anti-CTxB dilution at 1:100 (J). Despite this, no membrane bending was detected under these conditions, although the edge of the supported bilayer was readily apparent *(black area)* (E). Scale bar, 2 μm.

**Figure S3:**
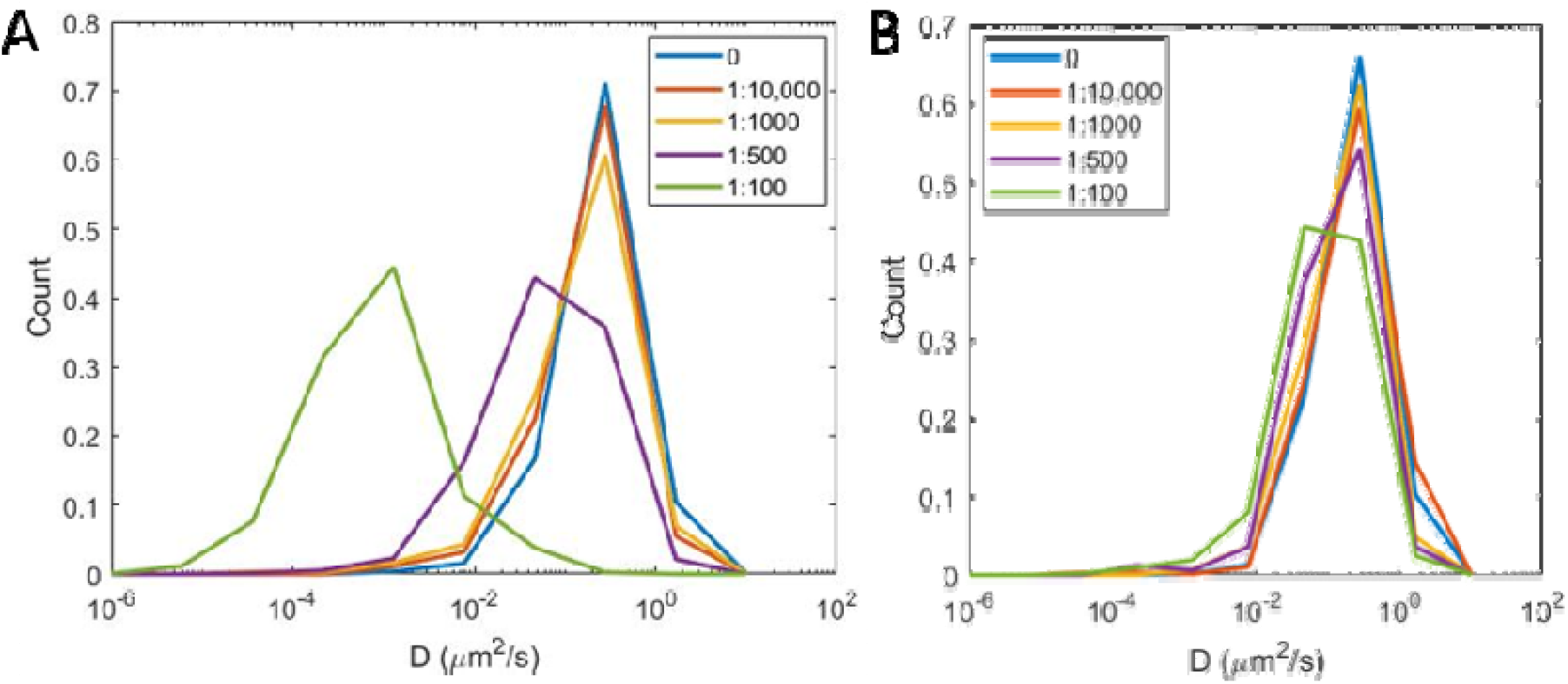
The diffusion of mCTxB, but not DiI, is slowed in the presence of increasing concentrations of anti-CTxB antibody. Histograms show the distribution of diffusion coefficients measured by single-particle tracking in planar supported lipid bilayers for mCTxB (A) or DiI (B) as a function of the indicated dilution of anti-CTxB antibody.

**Figure S4:**
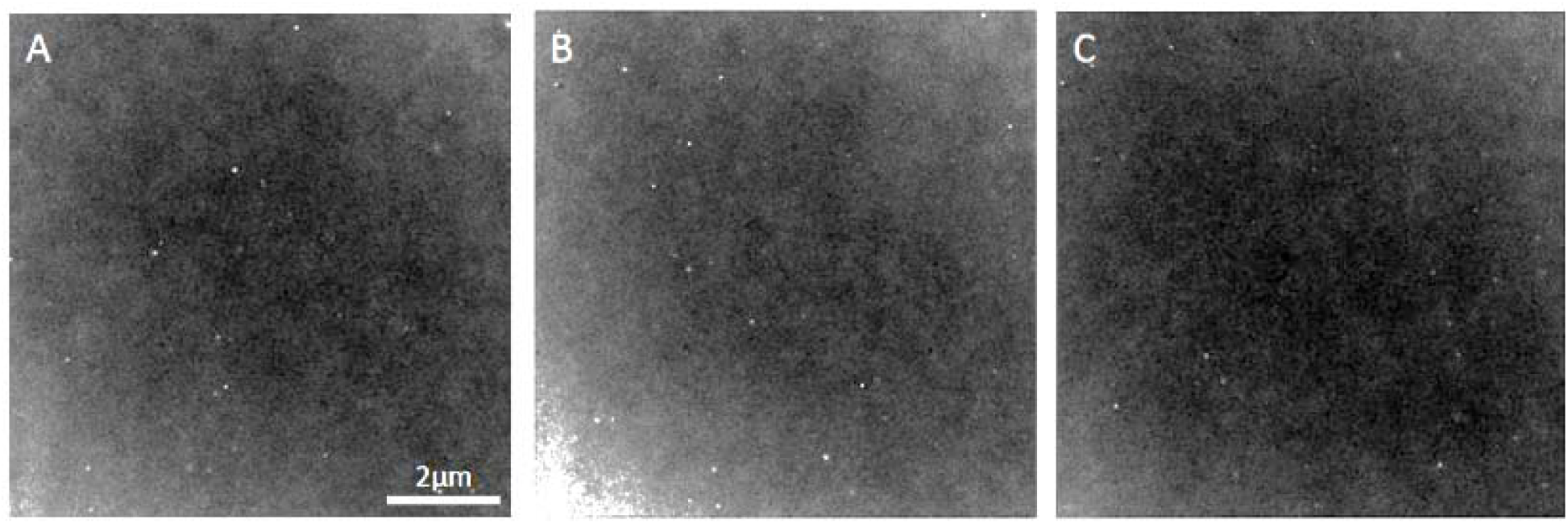
Generic crosslinking of lipids does not induce membrane budding. Planar POPC SLBs containing either (A) 0.3 mol%, (B) 1 mol%, or (C) 3 mol% DPPE-Biotin were labeled with 4.3 nM streptavidin 90 min prior to imaging via pPLM. Scale bar, 2 μm.

**Figure S5:**
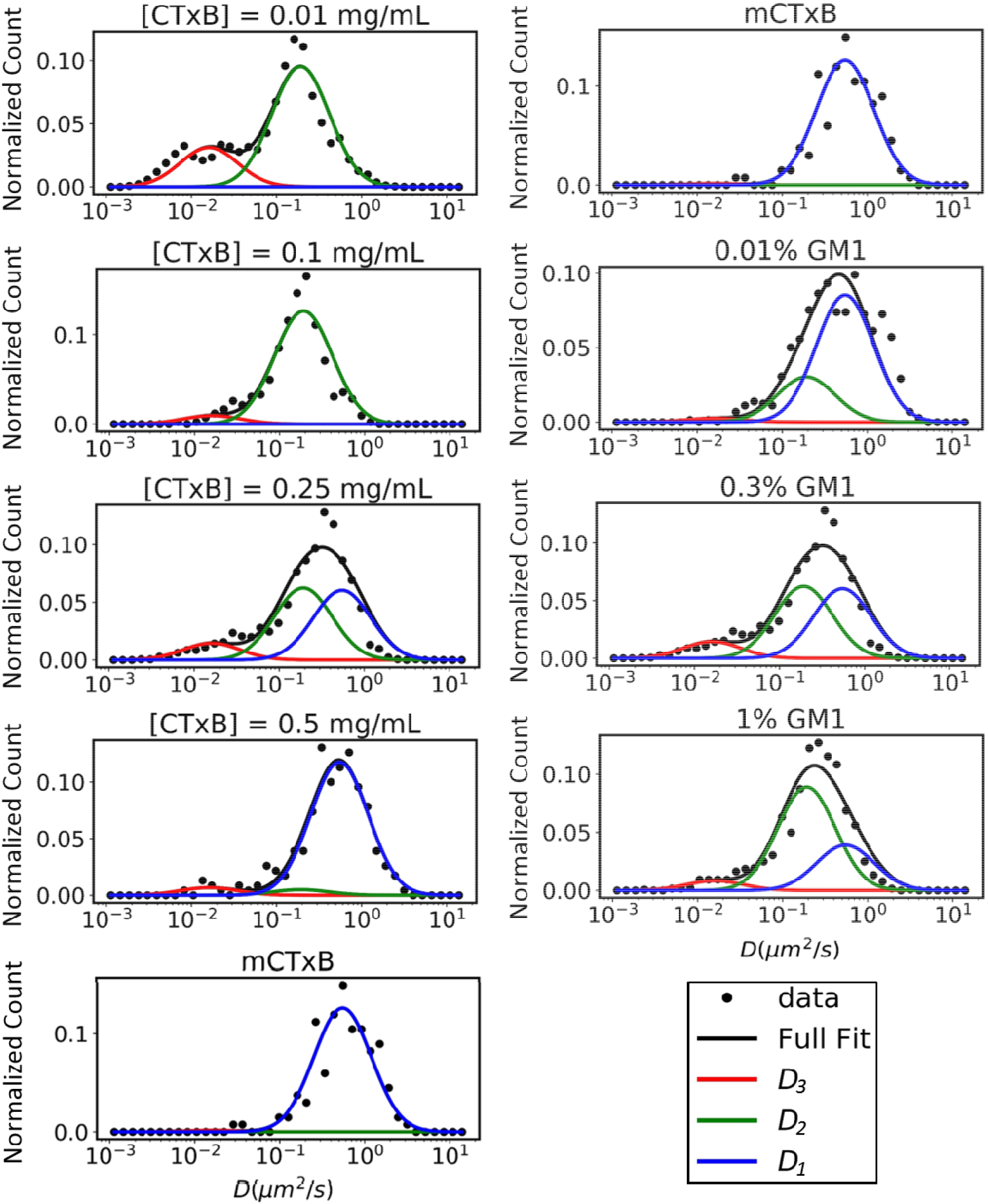
Fits to histograms of *D* for CTxB reveal three distinct subpopulations of diffusing species with different *D* values, whereas the histogram for mCTxB is consistent with the presence of a single population. Each histogram of *D* was fit to Eq. 1 (*black line*), allowing the amplitude of each fitting parameters to vary across experiments. The *D* values corresponding to each subpopulation (*D_1_*, *D_2_*, and *D_3_*) were held constant for all experimental fits. Note that the plot for mCTxB is shown in both columns to allow for better comparison across experimental conditions.

**Figure S6:**
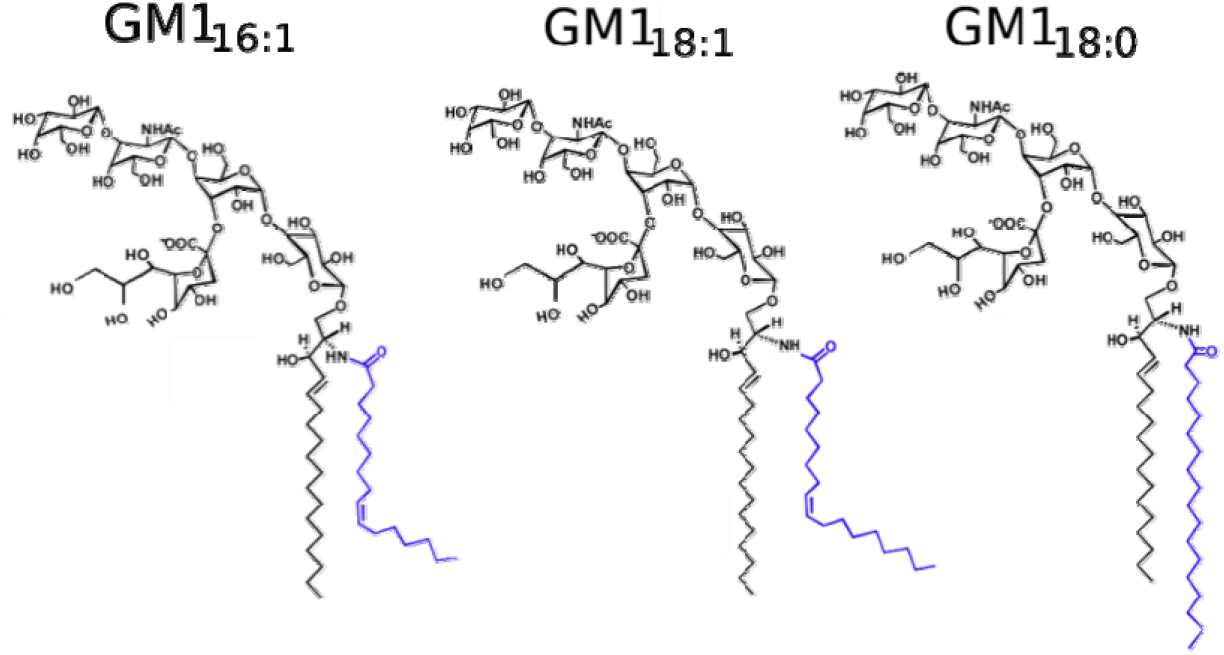
Chemical structure of custom-made GM1_16:1_, GM1_18:1_, and GM1_18:0_ (32). Note that commercially available ovine GM1 is predominantly composed of GM1_18:0_.

**Figure S7:**
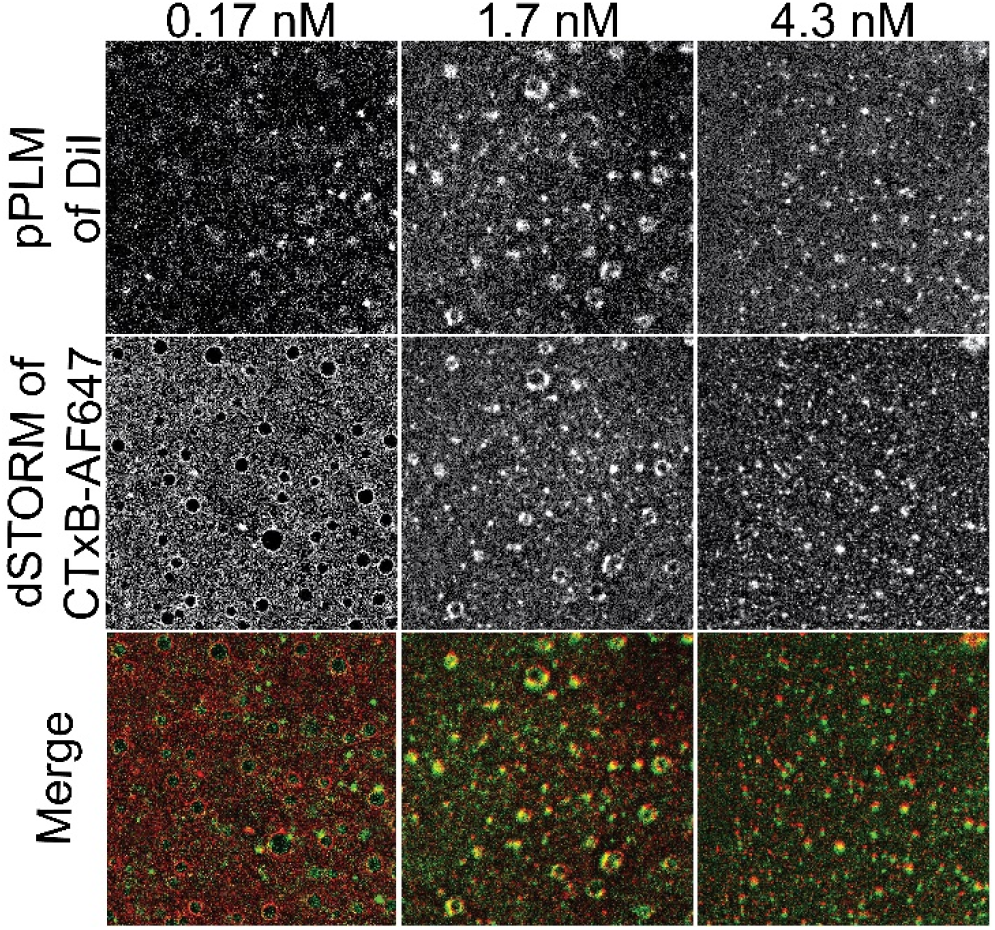
Increasing the ratio of GM1 to CTxB results in increased membrane curvature in model membranes containing cholesterol. The GM1:CTxB ratio was altered by labeling planar supported bilayers containing 0.3 mol% GM1 and 30 mol% cholesterol with either 0.17 nM, 1.7 nM, or 4.3 nM CTxB. Samples were then imaged by pPLM and dSTORM. Merged images show areas where clustered CTxB *(red)* colocalizes with sites of induced curvature *(green).* The presence of cholesterol had no significant effect on the membrane curvature created by CTxB. Scale bar, 2 μm.

**Figure S8:**
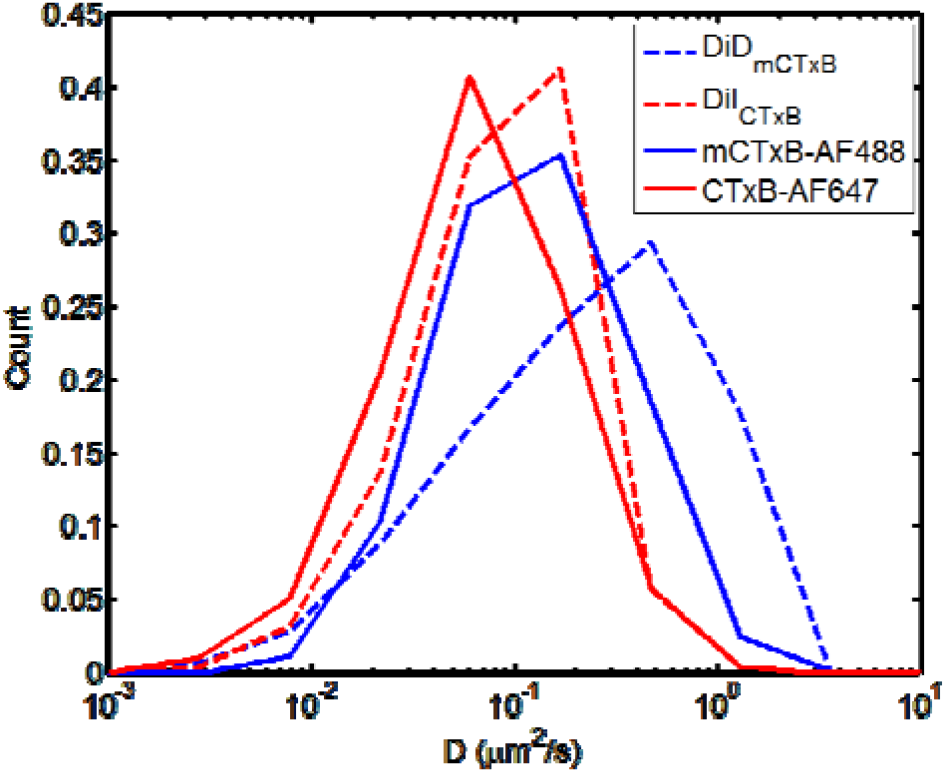
CTxB diffuses more slowly than mCTxB on live cells. Histograms of singlemolecule diffusion coefficients obtained in live COS-7 cells are shown for CTxB, mCTxB, and DiI or DiD in the presence of CTxB or mCTxB, respectively.

**Figure S9:**
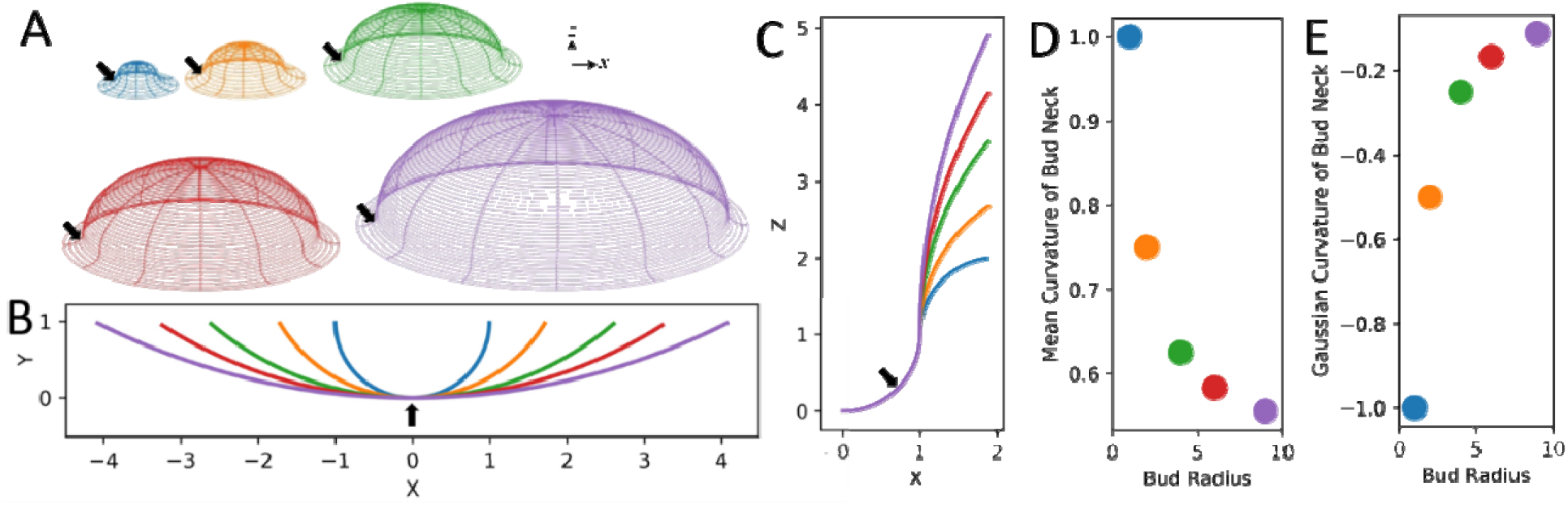
The necks of membrane buds provide one dimension of negative curvature. A model of the membrane buds of increasing radii are shown (A). The bud necks are highlighted *(black arrows).* The decreasing positive curvature in the X-Y plane (B) and consistently negative curvature in the X-Z plane (C) results in the mean curvature at the bud neck decreasing as the bud radius grows (D). In contrast, the Gaussian curvature increases to zero as a function of increasing bud radius (E). Additionally, a larger membrane bud would provide more membrane area on the neck to accommodate more CTxB.

